# Implicit adaptation is modulated by the relevance of feedback

**DOI:** 10.1101/2022.01.19.476924

**Authors:** Darius E. Parvin, Jonathan Tsay, Kristy V. Dang, Alissa R. Stover, Richard B. Ivry, J. Ryan Morehead

## Abstract

Given that informative and relevant feedback in the real world is often intertwined with distracting and irrelevant feedback, we asked how the relevancy of visual feedback impacts implicit sensorimotor adaptation. To tackle this question, we presented multiple cursors as visual feedback in a center-out reaching task and varied the task relevance of these cursors. In other words, participants were instructed to hit a target with a specific task-relevant cursor, while ignoring the other cursors. In Experiment 1, we found that reach aftereffects were attenuated by the mere presence of distracting cursors, compared to reach aftereffects in response to a single task-relevant cursor. The degree of attenuation did not depend on the position of the distracting cursors. In Experiment 2, we examined the interaction between task relevance and attention. Participants were asked to adapt to a task-relevant cursor/target pair, while ignoring the task-irrelevant cursor/target pair. Critically, we jittered the location of the relevant and irrelevant target in an uncorrelated manner, allowing us to index attention via how well participants tracked the position of target. We found that participants who were better at tracking the task-relevant target/cursor pair showed greater aftereffects, and interestingly, the same correlation applied to the task-irrelevant target/cursor pair. Together, these results highlight a novel role of task relevancy on modulating implicit adaptation, perhaps by giving greater attention to informative sources of feedback, increasing the saliency of the sensory prediction error.

## INTRODUCTION

Sensory feedback is continuously used to help keep that sensorimotor system calibrated, enabling the production of accurate movements despite ongoing changes to one’s body and environment. This adaptive recalibration process is thought to be driven in part or whole by sensory prediction error (SPE), the difference between the predicted and actual sensory feedback (Burge, Ernst, & Banks, 2008; Chaisanguanthum, Joshua, Medina, Bialek, & Lisberger, 2014; Kording, Tenenbaum, & Shadmehr, 2007; Mazzoni & Krakauer, 2006; Tsay, Haith, Ivry, & Kim, 2022; Wolpert, Ghahramani, & Jordan, 1995). In this paper, we examine what happens to sensorimotor adaptation when the sensory feedback is complex, ambiguous, and redundant.

When performing goal-oriented movements in the real world, the visual field is cluttered, possibly obscuring the feedback of the movement (Burge et al., 2008; Körding & Wolpert, 2004; Tsay, Avraham, et al., 2021; Tsay, Kim, Parvin, Stover, & Ivry, 2021; Wei & Körding, 2010) or even the target itself (Meegan & Tipper, 1998). Furthermore, the motion of other visual objects may be sources of distraction or interference. How does the motor system determine which signal is relevant for evaluating the consequences of an action, and how is this process influenced by the presence of competing visual signals?

Kasuga et al (2013) studied these questions using a center-out reaching task in which the participants could not see their moving arm, but had to rely on feedback provided by a moving cursor (Kasuga, Hirashima, & Nozaki, 2013). On most trials, a single cursor reflecting the participants true hand position was presented. Interleaved with these were perturbation trials. On some of the perturbation trials, the cursor was rotated relative to the true hand position by a variable angle, introducing a sensory prediction error. On other perturbation trials, there were two or three cursors, each following a different displaced trajectory. For example, one cursor might correspond to the true hand position with the other cursor(s) rotated by varying amounts; or all of the cursors might be rotated from the true hand position. Given that the number of cursors and their respective rotations was randomized across trials, implicit adaptation was measured by calculating the trial-by-trial change in hand angle. The results showed that the size of the trial-by-trial change could be modeled by taking the average predicted response to each of the individual cursors, albeit with a global attenuation in comparison to the single cursor condition. Thus, the response to two cursors, one at 0° (veridical feedback) and one rotated by 45° was similar to when there were two rotated cursors, one at 15° and one at 30°. The fact that there was adaptation even in the former case is especially surprising given that one cursor provided veridical feedback. The adaptation system did not appear to be preferentially sensitive to veridical feedback.

One issue of note in the Kasuga study is that, from the participants’ perspective, it may be appropriate to produce a composite error signal from the individual cursors because all of the cursors were potentially of equal importance and relevance to the task. However, in a natural environment, there is likely to be one relevant source of feedback amongst irrelevant and potentially distracting sources of information. To better understand how feedback drives sensorimotor adaptation, it is important to know whether it is sensitive to the relevance of available feedback. In the context of online feedback control, Reichenbach et al. (2014) provided a compelling demonstration that the motor system was indeed sensitive to the relevancy of visual signals (Reichenbach, Franklin, Zatka-Haas, & Diedrichsen, 2014). Participants performed reaching movements in which the feedback display included one cursor linked to the true hand position as well as up to 4 distractor cursors that moved with a similar but predetermined, and thus non-contingent velocity profile. At some point during the movement, one of the cursors made an abrupt lateral shift. Rapid, online corrections to the perturbation were much stronger when the perturbed cursor was the one linked to the hand position, compared to when the perturbed cursor was one of the distractors. Similar effects of task relevance have been observed in force-field adaptation studies (Heald, Ingram, Flanagan, & Wolpert, 2018), suggesting that implicit processes required for both on-line corrections and sensorimotor adaptation are sensitive to the task-relevance of different feedback signals.

In contrast, other lines of research have highlighted how sensorimotor adaptation is seemingly impervious to feedback regarding task goals and outcomes (Held & Gottlieb, 1958; Welch, 1969). Consider the aiming landmark task, first introduced by Mazzoni and Krakauer, (2006). After being briefly exposed to a 45° visuomotor rotation, the participants were instructed to aim to a landmark positioned 45° in the opposite direction from the target. By implementing this strategy, the participants were essentially perfect after one trial, producing movements in which the rotated cursor hit the target. Nonetheless, over the next 100 trials or so, performance deteriorated, with the hand angle increasing even further away from the target (Taylor & Ivry, 2011). This paradoxical result arises because the adaptation system, impervious to the strategy, recalibrates the sensorimotor mapping to reduce the SPE, here defined as the difference between where the movement was directed (towards the landmark) and where the cursor appeared (at the target). Similarly, participants implicitly adapt to the movement of a cursor that follows an invariant spatial trajectory displaced from the target, even when they are fully aware of the manipulation and told to ignore it (Avraham, Morehead, Kim, & Ivry, 2021; Kim, Morehead, Parvin, Moazzezi, & Ivry, 2018; Morehead, Taylor, Parvin, & Ivry, 2017; Parvin, McDougle, Taylor, & Ivry, 2018; Tsay, Parvin, & Ivry, 2020).

Experiments such as these have led to the view that implicit adaptation is solely dependent on SPE, impervious to feedback concerning the task outcome (Kim, Parvin, & Ivry, 2019). The results from the multiple cursor study of Kasuga et al. (2013) would also be consistent with this hypothesis. However, as noted above, the cursors were, in a sense, all task-relevant. Here we employ a multiple cursor task similar to Kasuga *et al*., but with two key changes. First, rather than randomize the perturbation from trial-to-trial, we employed a fixed rotation throughout the training period and assessed adaptation in a subsequent block where feedback was withheld. Second, and most importantly, we varied the cursor configurations and instructions as a way to manipulate the task relevance of the different visual feedback signals. In this way, we sought to determine whether implicit adaptation is sensitive to the relevance of the feedback.

## METHODS

### PARTICIPANTS

Undergraduate students (n = 64, 41 females, age = 20 ± 2 years) were recruited from the University of California, Berkeley community and financially compensated for their participation in the experiment. All participants were right handed, as assessed by the Edinburgh Handedness Inventory (Oldfield, 1971). The research protocol was approved by the UC Berkeley institutional review board.

### EXPERIMENTAL APPARATUS

The participant was seated in front of a horizontally oriented computer monitor that was supported by a table frame. All hand movements were tracked on a digitizing tablet (53.2 cm x 30 cm, ASUS), positioned 27 cm below the monitor. The participant held a modified air hockey ‘paddle’ embedded with a digitizing stylus to make center-out reaching movements over the tablet surface in response to visual stimuli displayed on the monitor. The participant’s hand was occluded by the table/monitor, and the room was minimally lit to further preclude visual feedback of the arm. The latency between the movement of the digitizing stylus and the updating of the cursor position on the monitor was 33 ms. The experimental code, controlling the visual display and acquisition of kinematic information was written in MATLAB (version 2016), using the Psychophysics toolbox extensions (Pelli, 1997).

### OVERVIEW OF THE REACHING TASK

Participants performed 8 cm reaches to targets located around a central starting location. The start location was indicated by a 6 mm white annulus, and the target was a 6 mm blue circle. The visual displays also included feedback cursors (3.5 mm white circles) that, depending on the condition, either corresponded to the participant’s hand position or were rotated around the start location at a fixed angle from the hand position.

In all conditions, participants were instructed to produce rapid movements such that the task-relevant designated cursor would ‘shoot’ through the target. Movement onset during the task was arbitrarily defined as the time at which movement amplitude reached 1 cm from the center of the start position. Movement time (MT) was defined as the duration from this point until the hand reached a radial distance of 8 cm, the target distance. Auditory feedback concerning MT was used to encourage participants to make relatively fast movements. For reaches shorter than 100 ms or longer than 300 ms, the messages ‘too fast’ or ‘too slow’ were played over the computer speaker. A neutral ‘knock’ sound was played if MT fell within the desired range. Across all trials, the median reaction time (RT) and MT were 462 ms and 172 ms, respectively. The median total trial time (TTT), defined as the time from the start of one trial to the start of the next, was 3714 ms. One way ANOVAs revealed no differences between groups for RT (*F_(_*_3, 44)_ = 0.448, *p* = 0.720), MT (*F*_(3, 44)_ = 1.336, p = 0.275), or TTT (*F*_(3, 44)_ = 0.245, p = 0.865).

On trials with visual feedback, the cursor or cursors were visible until the movement amplitude reached 8 cm, whereupon the end point position was frozen for an additional 1 s. By freezing the cursors, the participant received additional endpoint feedback of performance accuracy. At the end of the feedback period, the cursors were turned off and the participant moved their hand back to the start position. To help the participant find the start position, veridical feedback was provided when their hand was within 1 cm of the start position. Once in the start position, the feedback cursor was turned off and the annulus filled, indicating that the participant should prepare for the next trial. The next target appeared once the participant remained within the starting position for 500 ms.

### EXPERIMENT 1

Experiments 1a and 1b (n = 48, 12 per group) employed a similar design in which the participant completed a series of five blocks. The *No Feedback Baseline* block was composed of 24 reaches without visual feedback, one to each of 24 targets evenly spaced at 15° intervals (0° to 345°, with 0° corresponding to a rightward movement). This block was included to familiarize the participants with the experimental apparatus and with making movements in the desired time. The next block, *Feedback Baseline*, was composed of 10 cycles of reaches, with each cycle composed of one reach to each of the 24 target locations (240 trials). Veridical online feedback was provided by a feedback cursor aligned with the participants’ hand. Next was the *Training* block in which the specific experimental manipulations of the cursor feedback were introduced (detailed below). Targets were limited to three locations (30°, 150°, 270°) with 80 cycles of 3-target sets (Training, 240 total trials). The 120° spacing was chosen to minimize generalization/interference of adaptation effects between the three training locations (Day, Roemmich, Taylor, & Bastian, 2016; Krakauer, Pine, Ghilardi, & Ghez, 2000). The *Aftereffect* block had one cycle of 24 trials with participants reaching to each target without visual feedback, similar to *No Feedback Baseline*. At the start of this block, the participant was explicitly instructed to move their unseen hand directly to the target. The *Washout* block had the same structure but with veridical visual feedback (3 cycles, or 72 trials). The *Aftereffect* and provided the critical data to test for aftereffects, indicative of implicit adaptation and generalization. In all blocks, the order of the target location was randomized within a cycle.

#### EXPERIMENT 1a

Experiment 1a was designed to examine the influence of distractor cursors on performance during a visuomotor rotation task. We compared performance between training conditions in which the display contained a single feedback cursor rotated 45° from the true hand position or when the display also contained two additional cursors, positioned +/-45° relative to the single cursor (Fig 1D).

##### One Cursor group (n=12)

A single feedback cursor was visible during the blocks with visual feedback. In the *Feedback Baseline* and *Washout* blocks, the cursor provided veridical feedback. At the start of the Training block, the participant was informed that the feedback cursor would no longer be veridical but would now be displaced by 45° relative to their hand position (counterbalanced clockwise or counterclockwise across participants). They were instructed that the task goal was to compensate for this rotation such that the rotated cursor would hit the target. The experimenter explained the effect of the rotation on the cursor and the new task by illustrating it on a whiteboard. Although it could be inferred from the instructions that they could re-aim 45° away from the target to achieve the goal, we did not explicitly instruct them to use such a strategy. By making the rotation explicit, we believed there would be less ambiguity in the Aftereffect and Washout blocks, in which we assess implicit adaptation by instructing participants to reach directly with their hand. Furthermore, since participants were informed about the rotation of the cursors in the Three Cursor group (below), this made the number of cursors the only difference between the two groups.

##### Three Cursor group (n=12)

Three feedback cursors were visible during each of the blocks with visual feedback, with the cursors separated by 45°. Each cursor had a unique color: green, orange, and purple (RGB values (/255) [7 210 0], [231 145 53], [234 0 238] respectively). All colors were approximately matched on luminance based on a ‘Hue Chrome Luminance’ color scheme, and the assignment of color to cursor was counterbalanced across participants. In the *Feedback Baseline* and *Washout* blocks, the participant was informed that the middle cursor would correspond to the true hand position and instructed to hit the target with that cursor, specified in terms of the cursor color, idiosyncratic for each participant. This cursor was flanked by the two distractor cursors, resulting in three cursors moving at -45°, 0°, 45°, relative to hand position (Figure 1). The participant was instructed to ignore the other two cursors. Thus, if the color assignment for the -45°, 0°, 45° cursors was green, orange, and purple respectively, then the participant was to hit the target with the orange cursor and ignore the green and purple cursors.

During the Training block, all three cursors were rotated by 45°, such that the cursors now appeared at 0°, 45°, 90° (or 0°, -45°, -90°, counterbalanced across participants) relative to the true hand position. The participant was informed of the manipulation and instructed to hit the target with the middle cursor. Since the relative color assignment of the cursors remained the same, the instructions did not change. Using the color mapping example from above with a counterclockwise rotation, the participant’s goal was still to hit the target with the orange cursor (now rotated 45° from true hand position), while ignoring the green and purple cursors. With this arrangement, the purple cursor now ended up corresponding to the true hand position (0°) and the green cursor was rotated by 90°, relative to the hand. As such, if each cursor contributes equally to form a composite SPE (Kasuga et al., 2013), the net SPE in the Three Cursor condition is identical to that in the One Cursor condition.

#### EXPERIMENT 1b

As reported below, the inclusion of the two task-irrelevant distractors attenuated adaptation in the Three Cursor group, relative to the One Cursor group. Experiment 1b was designed to test two hypotheses that could account for this attenuation. The first hypothesis, “Error Averaging”, posits that adaptation is equally driven by SPE signals generated from all three cursors, but that their weightings add up to less than 1. Thus, the total amount of adaptation resulting from a 0°, 45°, and 90° would be less than from just one 45° cursor. The second, “Non-Specific”, hypothesis is that the presence of distractors dilutes the effects of adaptation in a general manner, and as such, the attenuation effect is not dependent on the particular path of the distractor cursors. To evaluate these hypotheses, we compared two, three-cursor variants in Experiment 1b, using the same trial structure as in Experiment 1a.

In the Compensate group (*n* = 12), the visual feedback was not rotated during the Training block. Instead, the participants were instructed to hit the target with the 45° side cursor. For example, if the participant was in the clockwise group (again, counterbalanced across participants) and had the color mapping shown in Figure 1D, they were instructed to hit the target with the green cursor. “Compensate” refers to the fact that, while the three cursors were not rotated relative to baseline, the instructions required participants to compensate for the angular offset of the task-relevant cursor. Note that as a result of this change in instructions, the angular offset of task-relevant cursor from the hand position (by 45°) is similar to the Three Cursor condition in Experiment 1a.

In the Ignore Rotation group (*n* = 12), the three cursors were rotated by 45° in the Training block. However, unlike Experiment 1a, the task goal was changed for this block, with the participant instructed to hit the target with the outer cursor that was in the opposite direction of the rotation (e.g., for a counterclockwise rotation, the task-relevant cursor now became the cursor that was clockwise to the center cursor). As shown in Fig. 1D, the net effect of the rotation and change in instructions results in the task-relevant cursor corresponding to the position of the hand. “Ignore” here refers to the fact that participants can ignore the 45° rotation applied to the three cursors since the task-relevant cursor ends up being veridical with respect to the participant’s hand.

In summary, the groups used in Experiment 1a and 1b match average and task-relevant sensory prediction errors across their conditions, offering a window into the unique contribution of these errors. Furthermore, with the exception of the Ignore condition, all groups were required to make a 45° change in reaching angle in the Training block to compensate for the perturbation, making reaching behavior very similar across these groups.

**Figure 1:**
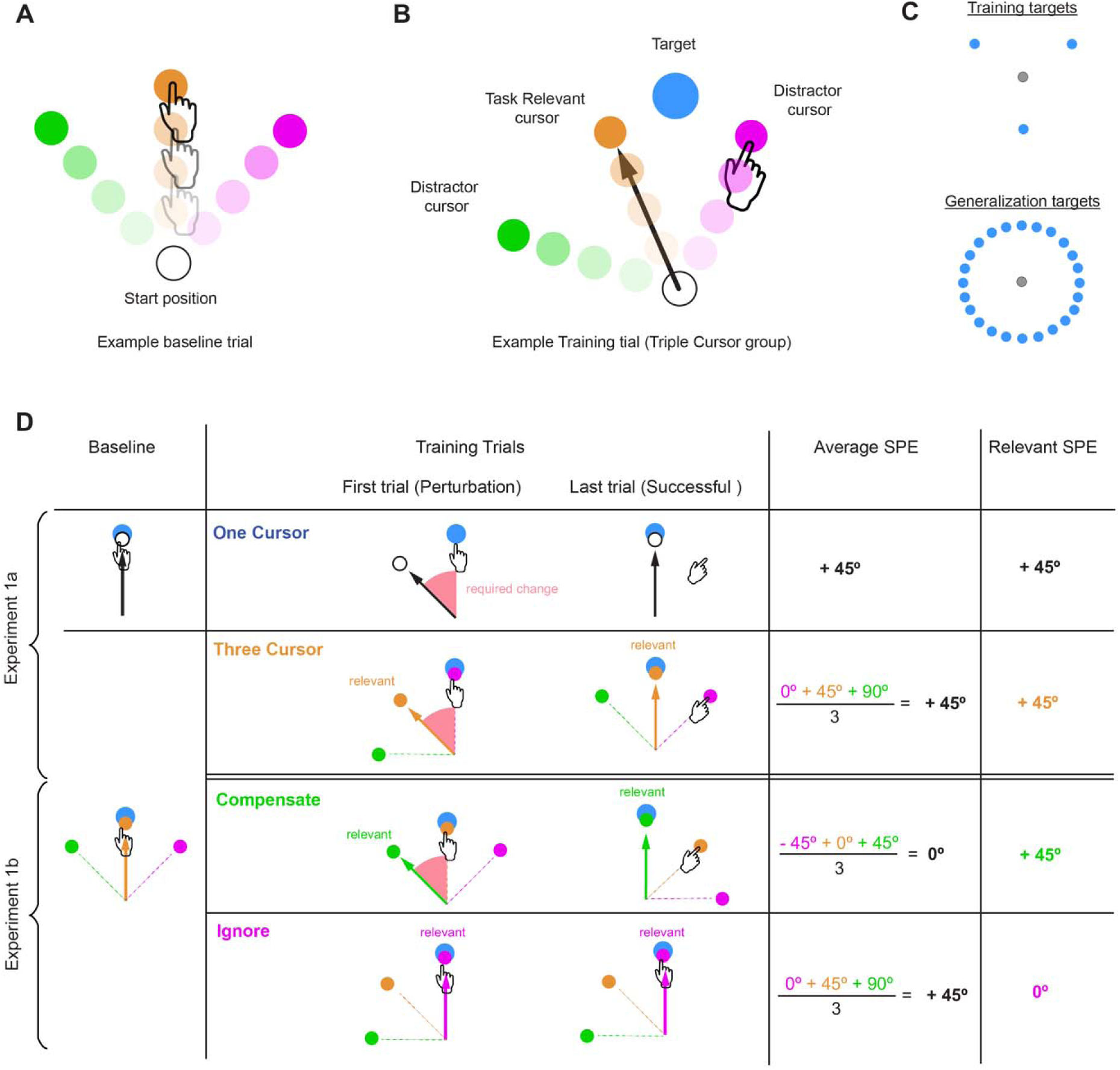
Feedback configurations for the Three Cursor group in Experiment 1. **A)** In the baseline block, the middle cursor follows the veridical hand position, with two additional cursors rotated either +/-45° on either side. Participants are told to reach directly to the target with the middle cursor. **B)** In the *Training* block, all the cursors are rotated by 45°, such that the middle cursor no longer corresponds to the true hand position. Participants are told to hit the target with the middle cursor and ignore the other two cursors. **C)** Target locations for Experiment 1. In the *Training* block, targets appeared in three locations (30°, 150°, 270°). In all other blocks, targets appeared in 24 generalized locations (from 0° to 345° in 15° intervals). **D)** Experimental conditions for Experiment 1. Experiment 1a compares learning from one cursor versus three cursors, keeping average SPE and relevant SPE the same. Experiment 1b employs two additional multiple cursor conditions to understand how the distractor cursors affect implicit adaptation. In the Compensate group, the three cursors were not rotated during the *Training* block, but the relevant cursor changed. Participants were instructed to hit with an outer cursor (green cursor), thus making that cursor relevant. This change in relevancy decreases the average SPE (i.e., 0°) but keeps the relevant SPE the same (i.e., 45°) as the Three Cursor group. In the Ignore group, the cursors were rotated by 45° during the *Training* block. Participants were told to hit the target with the cursor that followed their veridical hand position (pink cursor). Thus, the relevant cursor has change but the average SPE remains the same as the Three Cursor group.

### EXPERIMENT 2

Experiment 2 (*n* = 16) was designed to further investigate how adaptation is influenced by task-relevant feedback in the context of redundantly controlled objects. Generalization in this experiment was assessed in one half of the workspace, with reaches limited to 13 target locations spaced every 15° (Fig. 4C). To control for workspace-specific training artifacts (e.g., biomechanical biases), participants were assigned to reach in one of four areas of the workspace – Top (0°-180°), Left (90°-270°), Bottom (180°-360°), or Right (270°-90°). In the *No Feedback Baseline* block, the participant made three reaches to each of the 13 targets without visual feedback. This was followed by the *Feedback Baseline* block, consisting of five reaches to the 13 targets with veridical feedback provided by a single cursor.

In the *Training* block (200 trials, with a break at the midpoint), two targets were presented, positioned approximately +/-45° clockwise (see below) from the center of the participant’s workspace. Two cursors appeared during the reaching movement, one rotated clockwise (-45°) and the other counterclockwise (45°) from the true hand position. The participant was instructed that one target-cursor pair was task-relevant. For example, if the right target was task-relevant, then the participant’s goal was to hit this target with the cursor that was rotated in the clockwise direction. The participant was told to ignore the other target-cursor pair. The side of the task-relevant target was counterbalanced across participants. Note that by limiting the workspace to 180°, the instructions were stated in terms of a fixed direction (e.g., “hit the target on the right” or “hit the upper target”). Following the *Training* block, the participant completed an *Aftereffect* and *Washout* block (39 reaches each, three to each of 13 target locations). These blocks provided the data for assessing aftereffects and generalization.

To verify that the participant was attempting to hit the task-relevant target, the location of each target was independently jittered from trial to trial. The size of the jitter was one of five values, -10°, -5°, 0°, 5°, and 10°, and determined in a pseudorandom manner for each participant, such that each value was selected once every five trials. The overall order was pre-defined so that the correlation between the jitter for the two targets would be zero across the experiment. As a result of ensuring they were uncorrelated, the beta weights associated with each target were independent. By jittering the exact location of the target, we could perform a multiple regression, using each target as a predictor of hand angle. The beta weights obtained from this analysis indicate the participant’s sensitivity to the relevant and distractor target.

### DATA ANALYSIS

No statistical analyses were performed beforehand to determine sample size. The chosen sample sizes were based on similar studies of visuomotor adaptation (Huang et al., 2011; Galea et al., 2015; Vaswani et al., 2015), as well as considerations for counterbalancing. Experiment 1 (n=12 per group, 48 total) consisted of 2 participants for every cursor color combination (3) and direction (2, clockwise vs. counterclockwise). Experiment 2 (n= 16) had 2 participants in every possible counterbalanced configuration (4 workspaces x 2 directions).

The primary dependent variable of interest for all experiments was the heading angle of the hand. This was defined as the signed angular difference between the position of the hand and the target angle at peak radial velocity. Trials in which the hand angle was more than 90° away from the target were excluded from the analysis (42 total, 0.15% of all trials over all participants). Excluding these trials did not affect the outcome of any statistical tests. The data were averaged across cycles (24 successive reaches; 1 reach to each target), and baseline subtracted to aid visualization. Baseline was defined as mean hand angle over the last 3 movement cycles of the baseline phase with veridical feedback.

The degree of adaptation was quantified as the change in heading angle in the opposite direction of the rotation. We calculated heading angle during early adaptation, late adaption, and the aftereffect phase. Following previous studies (Kim et al., 2018; Tsay, Avraham, et al., 2021), early learning was defined as the mean hand angle over the cycles 3 – 7 to estimate the per trial rate of change during the *Training* block. (We also performed a secondary analysis using cycles 1–10 and obtained nearly identical results.). Late learning was defined as the mean hand angle over the last 10 cycles during the *Training* block. The aftereffect was operationalized as the mean hand angle over all cycles of the *Aftereffect* block.

All analyses were conducted using custom scripts in Matlab (version 2016). Our experimental design for Experiments 1a and 1b aimed to determine if there were group differences across adaptation blocks (early adaptation, late adaptation, and aftereffect blocks). Accordingly, we performed two-tailed t-tests to compare groups across these phases. We did not apply post-hoc corrections as our statistical tests were based on a priori hypotheses regarding expected group differences; nonetheless, the key results remain robust after applying Bonferroni corrections for the three comparisons. In Experiment 2, our design focused on the effect of target relevance. We quantified this by correlating heading angles with target angles; higher beta weights indicate greater attention to tracking the relevant/irrelevant jittered cursor/target location. The beta weights were evaluated with one-sample t-tests to determine if they differed from zero, and within-participant t-tests were used to compare beta weights across relevant and irrelevant targets. We report standard measures of effect size (Cohen’s d for between-participant comparisons; Cohen’s d_z_ for within-participant comparisons).

For the Gaussian fitting procedure in Experiments 1a and 1b, we used the ’fmincon’ function in MATLAB. We first fit the group average generalization curves using a Gaussian function characterized by three free parameters subject to the following lower and upper bound constraints: standard deviation [0°, 100°], height [0°, 100°], and mean [-100°, 100°]. To obtain 95% confidence interval estimates for these parameters, we employed a bootstrapping procedure 1000 times. For each iteration, we resampled N participants, where N represents the total number of participants in the experiment, with replacement. We then calculated the average generalization function from this bootstrapped sample and derived the three free parameters by fitting the Gaussian function.

For the Ignore group in Experiment 1b, we applied a cluster analysis approach to identify if there were any significant clusters during training where the hand deviated from 0 (Tsay et al., 2020). This step consisted of three steps. First, a t-test was performed for each cycle, asking if the observed hand angle diverged from zero. Second, clusters were defined as epochs in which the p value from the t-tests were less than 0.05 for at least two consecutive cycles. Third, to identify the probability of obtaining a cluster of consecutive cycles with significant p values, we performed a permutation test. In this, we created 1000 permutations of the data with the cycles shuffled. For each shuffled permutation, we performed the first two steps described above to identify clusters and for those meeting this criterion, we calculated the sum of the t-scores over the significant cycles. Doing this for each of the 1000 permutations resulted in a distribution of t-scores. The proportion of random permutations which resulted in a t-score of equal or greater to that obtained from the data could therefore be directly interpreted as the p value.

Applying the first two steps to the actual data, we identified only one cluster, and only of length two (training cycle 10 mean = 2.32° [1.29°, 3.35°], t(11) = 4.956, p = 4.315e-4, d = 1.431, and cycle 11 mean = 1.58° [0.55°, 2.62°], t(11) = 3.362, p = 0.006, d = 0.971). The sum of t-scores for the two consecutive cycles (t = 8.318) was compared to the distribution generated by the permuted distribution, producing a p-value of 0.004. We therefore concluded that this cluster of cycles represented a significant deviation from 0, rather than being due to chance.

### DATA AND CODE AVAILABILITY STATEMENT

Raw data and analysis code can be openly accessed at https://github.com/DariusParvin/Adaptation_Multiple_Cursor_Experiments.

## RESULTS

### EXPERIMENT 1a

In Experiment 1, we examined the impact of task-irrelevant cursors on implicit adaptation. We first compared adaptation to displays consisting of either a single cursor (One Cursor group) or three cursors, separated by 45° (Three Cursor group, Fig 2). In both groups, participants were informed that, during the training block, the task-relevant cursor (single or middle) was rotated by 45° relative to their hand position, and that their task was to hit the target with this cursor. In the Three Cursor group, participants were also told that one of the distractor cursors would coincide with their veridical hand position but that they should ignore it and focus on the task-relevant cursor.

Performance of the two groups during the Training block was compared to assess effects of the distractor cursors on overall learning. Both groups appeared to compensate for the perturbation at a similar rate and extent. There were no significant differences in early learning (cycles 3-7, One Cursor mean hand angle = 18.00° [9.90°, 26.10°], Three Cursor mean = 26.67° [17.23°, 36.12°], [*t*_(22)_ = -1.535, *p =* 0.139, d = -0.626]), nor in late learning (last 10 cycles, One Cursor mean = 43.44° [41.27°, 45.62°], Three Cursor mean = 41.55° [40.14°, 42.96°], [*t*_(22)_ = 1.608, *p =* 0.122, d = 0.656]). Previous studies have shown that implicit adaptation is typically limited to 10°-25° of learning (Bond & Taylor, 2015; Kim et al., 2018; Morehead et al., 2017; Tsay, Lee, Ivry, & Avraham, 2021). Given that the change in hand angle is much larger than this range and that the participants were explicitly informed of the manipulation, we assume there is a strategic (Tsay et al., 2023), aiming contribution to performance here. As such, the results indicate that the distractor cursors did not have an appreciable influence on the participants’ ability to adopt an aiming strategy to complement adaptation.

To assess implicit adaptation, we included a block of trials after training in which the cursor was no longer presented (Aftereffect block), and participants were asked to forgo strategy use and aim directly to the target (i.e., “Move your hand straight to the target as fast and accurately as you can. Do not aim away from the target”). Both groups exhibited aftereffects at the trained target location. However, the magnitude of adaptation differed for the two groups: The One Cursor group had a significantly greater aftereffect than the Three Cursor group (One Cursor group = 18.10° [14.50°, 21.69°], Three Cursor group = 9.72° [6.48°, 12.96°], [*t*_(22)_ = 3.808, p 9.622e-4, d = 1.555]).

There are at least two reasons why the aftereffect would be larger in the One Cursor group. First, the difference could reflect that adaptation is attenuated by the inclusion of task-irrelevant distractors. Alternatively, there could be a group difference in the use of an aiming strategy. Assuming that the generalization function for adaptation is centered on the aiming location (Day et al., 2016), differences in strategy use could cause differences in measured adaptation at the training location on aftereffect trials.

To evaluate these two hypotheses, we assessed generalization during the aftereffect block by measuring adaptation across a set of probe targets that spanned the workspace in 15° increments (Figure 1C). The attenuation hypothesis predicts that the functions would be aligned but with a lower peak for the Three Cursor group; the aiming hypothesis predicts that the functions would be similar in amplitude but misaligned. The results were consistent with the attenuation hypothesis. While the generalization functions for both groups were shifted towards the presumed aiming direction of aiming, the peak of the generalization function was similar for the two groups. Based on parameters obtained when fitting a Gaussian curve to the group data, the peak of the generalization function was 7.7° [4.4, 10.9] and 9.0° [5.1, 12.8], with considerable overlap of the confidence intervals for the One Cursor and Three Cursor groups, respectively [mean, bootstrapped 95% CI]). In contrast, the heights of the peaks were different for the two groups (One Cursor: 18.8° [16.7, 20.8]; Three Cursor: 11.0° [9.3 12.2]). These findings suggest that the presence of the additional cursors attenuated the magnitude of implicit adaptation.

#### Error Averaging vs Relevant error models of adaptation

We next asked how the inclusion of the distractor cursors in the Three Cursor group influenced adaptation. Specifically, we formulated two possible models to account for the attenuation.

The Error Averaging model, inspired by Kasuga and colleagues (Kasuga et al., 2013), proposes that the error from each cursor is processed simultaneously and contributes towards learning (Equation 1). By this model, the attenuation effect for the Three Cursor group is reflected in a reduction in the learning term, B_t_, relative to what it would be in the One Cursor case. Kasuga and colleagues observed a reduction of 0.57 for B_t_ in their three cursor conditions relative to the One Cursor condition. This value is consistent with our observed attenuation of 0.55 for the Three Cursor group in Experiment 1a (Three Cursor aftereffect divided by the One Cursor aftereffect).

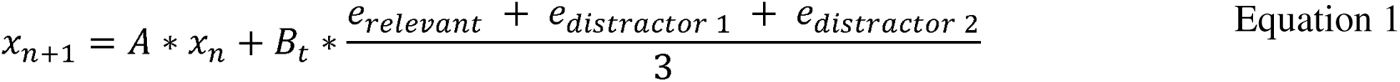

An alternative model, the Relevant Error model, proposes that implicit adaptation learns selectively from the relevant error, while the distractor cursors attenuate adaptation in a non-specific fashion. Thus, the directions of the distractor cursors have no bearing on the direction or magnitude of the aftereffect. The attenuation due to the presence of the two distractor cursors would also be manifest in a reduced B_t_ term, just as in the Error Averaging model (Equation 2).

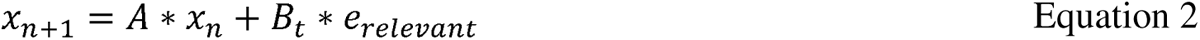

**Figure 2.**
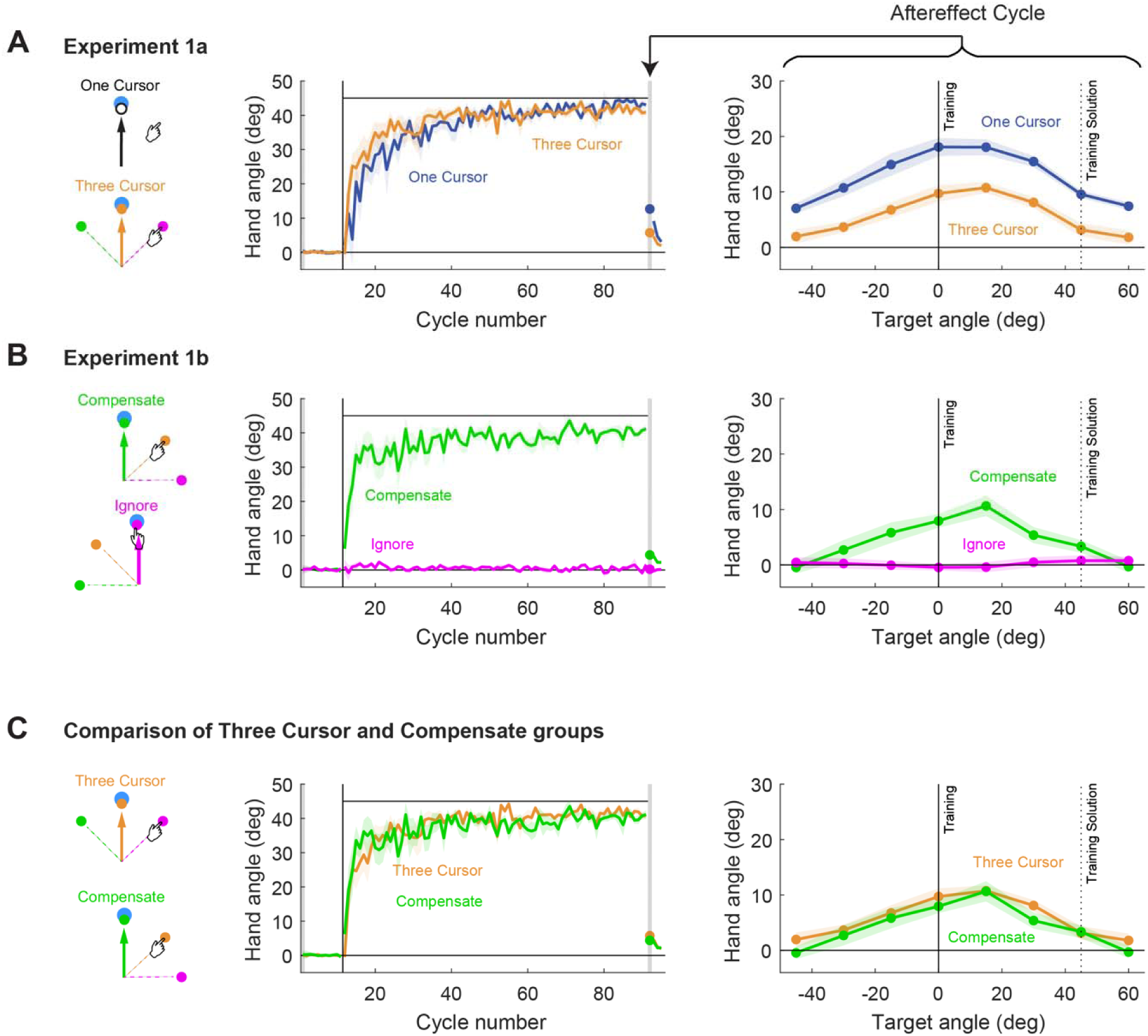
Experiment 1 Results. **A)** One Cursor and Three Cursor conditions in Experiment 1a. Middle: Both groups learn and compensate for the rotation at the same rate during training. Right: After the *Training* block, participants were instructed to reach directly to the generalization targets; no feedback was provided. The peak location of the generalization curves did not significantly differ, but the peak amplitude was greater in the One Cursor group compared to the Three Cursor group. **B)** Compensate and Ignore conditions in Experiment 1b. Middle: No change in hand angle was observed in the Ignore group. Right: Significant aftereffects were only seen in the Compensate group. **C)** Direct comparison of the Three Cursor and Compensate groups. No significant differences were observed in either the training (middle) or aftereffect (right) periods. Thick lines denote group means. Shaded regions denote +/-SEM.

**Figure 3.**
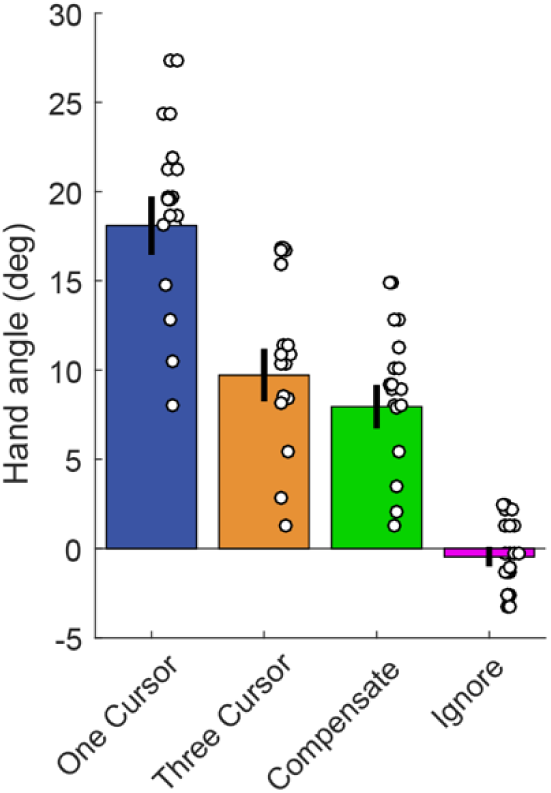
Results from the aftereffect block in Experiment 1. Three Cursor and Compensate groups have similar levels of aftereffects. Both groups also had significantly less than the One Cursor group, and significantly greater aftereffects than the Ignore group. Error bars denote +/-SEM. Dots denote individual mean aftereffects.

### EXPERIMENT 1b

To compare these two models, we tested two additional groups in Experiment 1b. The difference between the two models is in their treatment of the distractor cursors. In the Error Averaging model, feedback information from each cursor is equally weighted to form a composite error signal, whereas in the Relevant Error model, only the feedback information from the task-relevant cursor is used to define the error signal. Given that Error Averaging and Relevant Error models make identical predictions for the Three Cursor group, we devised two, three-cursor variants that yield divergent predictions for the two models.

In the Compensate group (Figure 2), the mapping between hand position and cursor is not changed in the Training block; that is, the cursor that provided veridical feedback during the initial Feedback block continues to provide veridical feedback in the Training block. However, the instructions change, with the task goal now requiring the participant to hit the target with the cursor that is offset 45° from the hand (45° clockwise or counterclockwise, counterbalanced across participants). To achieve this, the participant must move in the opposite direction (e.g., clockwise if the task goal is to his target with a counterclockwise cursor). While the required hand trajectory is the same as in the Three Cursor group, the set of errors is different. For the Compensate group, the sum is now 0° (-45°, 0°, 45°, relative to the hand direction). Thus, the Error Averaging model would predict no adaptation. In contrast, the Relevant Error model would predict the same amount of adaptation as observed with the Three Cursor group. Even though the errors from the distractor cursors are different between the Compensate and Three Cursor groups, both have a task-relevant cursor that is offset by 45° from the hand direction.

Consistent with the Relevant Error prediction, participants in the Compensate group exhibited a robust aftereffect with Gaussian shaped generalization (mean at training location = 7.95° [5.26°, 10.63°]; one sample t-test against 0: *t*_(11)_ = 6.510, *p* = 4.368e-5, d = 1.879). Furthermore, this group behaved similarly to the Three Cursor group during both the Training and Aftereffect blocks. There were no significant differences during the Training block in their early learning (mean and 95% CI, Three Cursor = 26.67° [17.22°, 36.11°], Compensate = 32.09° [25.70°, 38.48°], [*t*_(22)_ = -1.047, *p =* 0.306, d = -0.428]), nor in their late learning (mean and 95% CI, Three Cursor = 41.55° [40.14°, 42.96°], Compensate = 40.37° [38.34°, 42.40°], [*t*_(22)_ = 1.053, *p =* 0.304, d = 0.430]). There were also no significant differences in their aftereffects at the training locations (Three Cursor mean = 9.72° [6.48°, 12.96°], Compensate mean = 7.95° [5.26°, 10.63°], *t_(22)_* = 0.925, *p* = 0.365).

The Ignore group provided a second test of the models. Here, a rotation of all three cursors was introduced during the Training block, identical to that used for the Three Cursor group. However, the participants were instructed in this block to hit the target with the 0° cursor, the one that was now veridical with hand position. “Ignore” refers to the fact that the participant’s task was essentially to ignore the rotation by focusing on the (new) cursor that matched their hand position. The Error Averaging model predicts the same aftereffect as the Three Cursor group, since it is insensitive to the relevance of the error signals and the average error is again 45° (90°, 45°, 0°, relative to the hand direction). The Relevant Error model, on the other hand, predicts no aftereffect, since the hand direction and task-relevant cursor are aligned. Again, the results of the aftereffects conformed to the Relevant Error prediction. A t-test showed no significant aftereffect at the training location (mean = -0.45° [-1.66°, 0.75°]; one sample t-test against 0: *t_(11)_* = -0.833, *p* = 0.423, d = -0.240).

The training period of the Ignore group lent itself to testing another prediction of the Error Averaging hypothesis. If a composite SPE was present during the training trials, we should observe a ‘drift’ away from the target, despite initial accurate performance (Mazzoni, 2006; Taylor and Ivry, 2011). A t-test against 0 produced a non-significant result (mean = 0.57° [-0.20°, 1.33°], *t*_(11)_ = 1.635, *p* = 0.130, d = 0.472). We were worried, however, that this measure would not be sufficiently sensitive to capture what might be a transient effect (Taylor and Ivry, 2011). As a more sensitive alternative, we opted for a cluster analysis, assessing if there were any consecutive cycles with significant drift. This approach identified a significant cluster of two consecutive cycles (training cycles 10 and 11) in which the mean hand angle was greater than 0° (mean = 1.95° [1.09°, 2.81°], *t*_(11)_ = 4.992, *p* = 4.076e-4, d = 1.441). While this small cluster is in the expected direction, it is very small and of shorter duration than that observed in previous studies (e.g., a 15° drift that lasts for about 80-100 trials, see Taylor & Ivry [2011]). Thus, if there was any implicit adaptation in the ignore group, it was very limited.

Overall, the results rule out the Error Averaging model and are consistent with the Relevant Error model: Significant aftereffects were observed when the task-relevant cursor was offset 45° from the hand position during the Training block, either because we imposed a perturbation or altered the instructions. Moreover, the results from the Ignore group, as well as the similarities between the Compensate and the Three Cursor group, suggest there is minimal, or no specific influence from the distractor cursors. Rather, the presence of the distractors appears to have a non-specific attenuation effect on adaptation to the task-relevant cursor.

### EXPERIMENT 2

Although the results of Experiment 1 suggest that task relevance modulates adaptation, they are far from conclusive in this regard. Not only was there no imposed rotation for the Ignore group, just a change in the task-relevant color in the Training block, this group also did not have to aim away from the target (as in the Compensate group). To provide a more direct assessment of the contribution of task-irrelevant information, we conducted a second experiment, using a design in which there were two targets positioned approximately 90° apart, and two associated cursors, one rotated 45° in the clockwise direction from the hand direction and the other rotated 45° in the counterclockwise direction (Figure 4A). One target/cursor was designated task-relevant, and the other was designated a distractor (i.e., task-irrelevant). Regardless of which was task-relevant, the movement required to hit the target was directed around the midpoint of the two targets, resulting in each cursor landing near its associated target. We measured the aftereffects around each target location to determine the degree of adaptation. We hypothesized that the task-relevant target may form the locus of generalization for implicit adaptation. Therefore, whether there is also an aftereffect around the distractor target was of primary interest.

To verify that participants were following the instructions, we jittered the exact position of the targets (see Methods) and calculated the trial-by-trial hand angle relative to each target (Figure 4D). Using these time series, we then performed a multiple regression using the positions of the relevant and distractor targets on each trial to predict the hand angle. This analysis produced a beta weight for each target, quantifying how much the participant’s behavior reflected the tracking of each target (Figure 4E). We saw that greater weight was given to the task-relevant target in 15 of the 16 participants. The other participant appeared to not follow directions (beta weights were -0.14 and 0.09 for the relevant and distractor targets), and this person’s data were excluded from the remaining analyses.

The mean beta weight for the task-relevant target was 0.76 [0.67, 0.84] and for the distractor target was 0.20 [0.09, 0.32]. These values were significantly different from one another (*t_(28)_* = 8.149, *p* = 7.16e-9, d = 2.975). Interestingly, the beta weight for the task distractor target was significantly greater than zero (*t_(14)_*= 3.675, *p* = 0.002, d = 0.949). Thus, while the participants followed the instruction to track the relevant target, the results indicate that the participants were not able to completely ignore the distractor target. One possibility is that the distractor target may have served as an additional visual reference for aiming.

To assess adaptation, participants made reaching movements without feedback to an array of targets, spanning the workspace that encompassed the two positions of the targets during the training block. As in Experiment 1, the participants were instructed to reach directly towards each target. To analyze these data, we collapsed across the two groups (clockwise or counterclockwise rotation of the task-relevant cursor), displaying the data as though all participants were instructed to hit the clockwise target and ignore the counterclockwise target (Figure 4B). A significant aftereffect in the direction consistent with the rotation was observed around the task-relevant target location (mean 10.87° [6.48°, 15.27°]; one sample t-test against 0: *t_(14)_* = 5.304, *p* = 1.114e-4, d = 1.370). Interestingly, there was also a significant aftereffect at the distractor target location (mean -4.37° [-8.27°, -0.47°], *t_(14)_*= -2.406, *p* = 0.031, d = -0.621). The negative aftereffect associated with the distractor target was consistent with the distractor cursor having the opposite rotation to the relevant cursor. When we directly compared the aftereffects across the two targets by flipping the sign of distractor cursor, aftereffects were marginally greater in the task-relevant target compared to the distractor cursor (*t_(14)_* = 2.00, *p* = 0.067) with a medium effect size (d = 0.51).

The fact that we observed a small aftereffect around the distractor target location would seem at odds with the results of Experiment 1 where the effect of the distractor cursors was not dependent on their direction (e.g., non-specific attenuation). However, as shown in the beta weight analysis, the participants in Experiment 2 did not completely ignore the distractor target. To look at the relationship between tracking and adaptation, we examined the correlation between beta weights and magnitude of the aftereffect for both the relevant and distractor locations. To capture how much a participant weighted the relevant target over the distractor, we normalized the beta weights (beta weight for relevant target / sum of both beta weights) in these correlations. For the relevant location, the correlation was significant (*r* = 0.67 *p* = 0.007), meaning that those who were most successful in tracking the relevant target had the largest aftereffect around that target’s associated location. We performed the same analysis for the distractor target. Since the distractor cursor had the opposite rotation, the predicted correlation would be negative. This analysis revealed a similar pattern, but the correlation was not significant (r = -0.30, p = 0.274). We note that this analysis may be underpowered, given that the range both aftereffect and beta weights are narrower for the distractor location compared to the relevant location.

**Figure 4.**
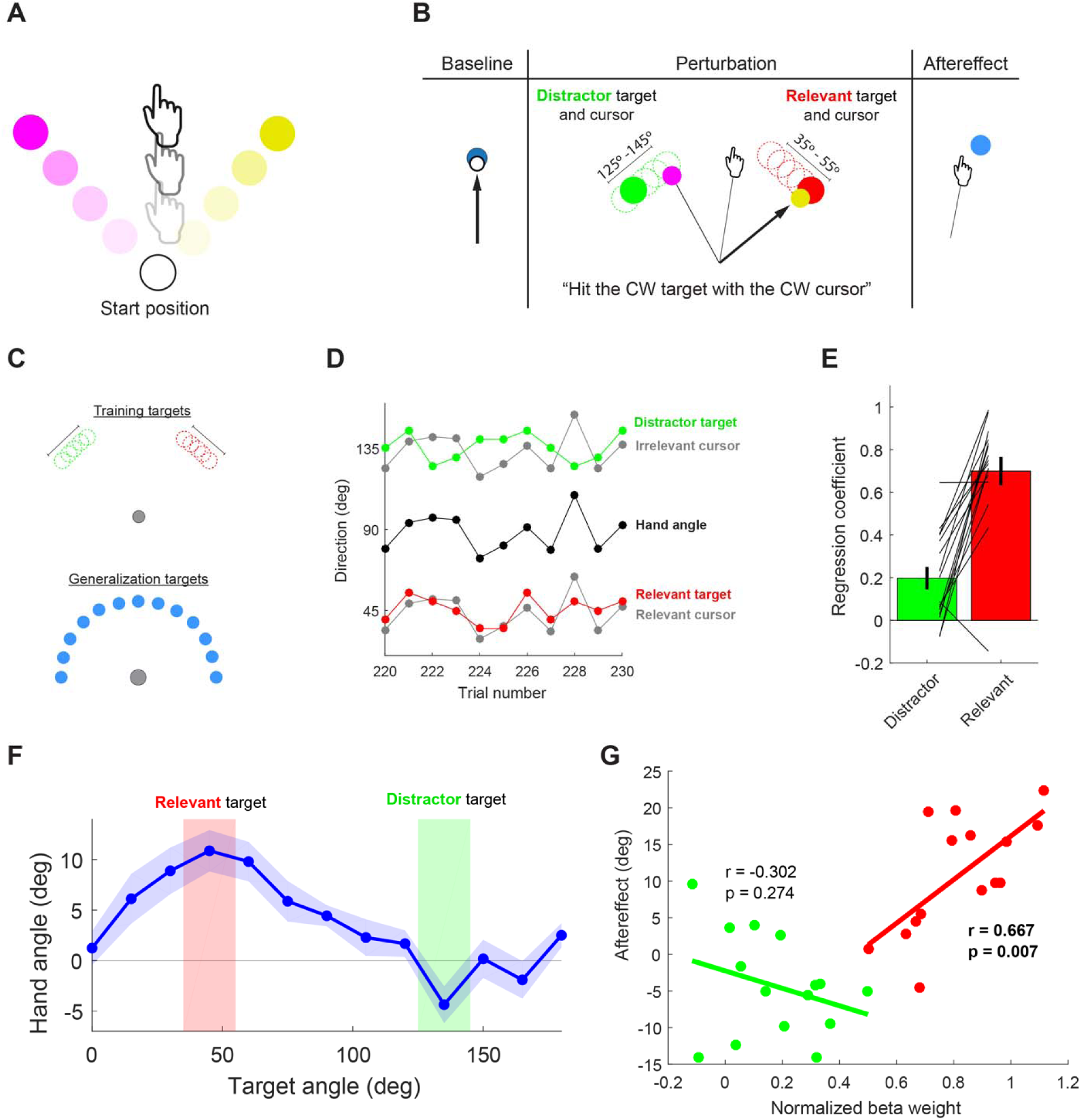
Results of Experiment 2. **A)** Feedback configuration for training trials. Two cursors appeared during the reach, rotated +/-45° to the veridical hand position. **B)** Experimental task. Baseline and aftereffect trials had one-cursor feedback reaches to generalization targets. During training, one target/cursor pair was made task-relevant by instructing the participant to focus on hitting that target. By jittering the targets, we can assess how well the participant tracked the relevant target. **C)** Target locations, limited to one half of the workspace. Top: Training target locations, centered +/-45° from the midpoint of the workspace. The exact location of each target was independently jittered by one of 5 values. Bottom, 13 generalization targets spaced 15° apart. **D)** Trial by trial tracking behavior for distractor (green) and relevant (red) target/cursor pairs. **E)** Beta weights for the relevant target (red) are significantly greater than for the distractor target (green). Data are group mean +/-SEM with lines representing individual performance. **F)** Generalization functions. The magnitude of the aftereffect is larger around the relevant target location (red) compared to the irrelevant distractor location (green), although the latter is also significantly greater than zero. G) Correlation between beta weights and aftereffect for relevant and distractor targets. Significant correlation for the relevant target (red) shows that participants who were better at tracking also had larger aftereffects. Non-significant correlation for distractor target (green). Note that in Panels D and F, the data are transformed to graphically depict the relevant target at 45° for all participants.

## DISCUSSION

In this study, we asked how implicit adaptation, an important process for maintaining calibration within the sensorimotor system, is affected by the presence of multiple visual signals. Through our use of task instructions, we varied the information value of the signals, designating one cursor as task-relevant and the others as task-irrelevant. As expected, we found that adaptation was sensitive to the rotation of the task-relevant cursor. Moreover, as shown in Experiment 2, the degree to which participants tracked the target with the relevant cursor predicted the size of their aftereffect, pointing to a strong relationship between the task relevance assigned to the cursor and the amount of adaptation. Interestingly, implicit adaptation was attenuated by the presence of distracting feedback, but this effect seemed to be non-specific: The presence of the distractor cursors reduced the magnitude of adaptation, but not the direction of adaptation. These findings highlight a novel role of task relevance for implicit adaptation.

By presenting multiple cursors and manipulating their relevance, we were able to make sense of two seemingly contradictory findings in the sensorimotor adaptation literature. The results from several studies suggest the motor system is sensitive to the relevance of feedback (Heald et al., 2018; Reichenbach et al., 2014). In contrast, the results from other studies have shown that implicit adaptation is insensitive to task goals. For example, when only one cursor is presented, implicit adaptation will learn from the SPE, regardless of whether participants are explicitly told to ignore the visual feedback, or even when the response to that cursor is detrimental to performance (Mazzoni & Krakauer, 2006; Morehead et al., 2017; Taylor & Ivry, 2011; Taylor, Klemfuss, & Ivry, 2010). Our results suggest a hybrid position: When more than one source of visual motion feedback is present, the primary input to the adaptation system is the most relevant source of visual motion feedback. This selectivity constraint has been observed in studies of other implicit motor functions, such as the finding that online corrections are faster in response to perturbations of a task-relevant cursor (Reichenbach et al., 2014) or that separate force fields can be learned depending on what part of a virtual tool is deemed relevant (Heald et al., 2018).

Although adaptation was sensitive to the relevance of the feedback, the overall attenuation of learning demonstrated that the selectivity was imperfect. The presence of multiple cursors had a similar attenuating effect in the present study as observed in Kasuga et al. (2013). Specifically, in both studies, adaptation was attenuated by about 45% when there were three cursors relative to a standard single cursor condition. This similarity is striking given that Kasuga used unpredictable rotations, interleaved the single and multi-cursor conditions, provided no instructions concerning relevance, and measured learning using a trial-by-trial, whereas we used a predictable rotation, blocked the conditions, instructed the participants to attend one one cursor, and measured learning in an aftereffect block. An attenuation of motor responses due to the presence of irrelevant feedback has also been observed for online corrections to cursor jumps (Reichenbach et al., 2014).

The Relevance Model posits that the attenuation is non-specific; it could be that the irrelevant information diverts visual attention away from the relevant cursor, thereby reducing adaptation. Previously, Taylor and Thoroughman (2007) demonstrated that participants adapted less to perturbations in a force field reaching adaptation task when they were engaged in a secondary task designed to divide their attention (Taylor & Thoroughman, 2007). Given the similarity between the cursors, namely that all originated from the same start position and moved with the same spatiotemporal correlation, it is likely that the irrelevant cursors resulted in some degree of attentional diversion in our task (Folk, Remington, & Johnston, 1992). This attentional explanation is also relevant when considering why participants are sensitive to task-relevance in the presence of multiple sources of feedback yet are unable to ignore a single cursor when instructed to do so.

However, the effect of visual distractors may be more nuanced than the non-specific attenuation we have posited. Specifically, in Experiment 1b, the visual distractors in the Ignore condition elicited negligible implicit adaptation, whereas in Experiment 2, the visual distractors elicited significant aftereffects around the irrelevant target locations. We attribute this discrepancy to at least two factors:

First, the presence of a veridical cursor and its associated target may modulate the degree to which the system adapts to irrelevant visual errors. In the Ignore condition of Experiment 1b, where both the veridical cursor and an aiming target are present, participants may fully attend to the veridical cursor/target pair, thus eliminating implicit adaptation in response to the irrelevant visual errors. In Experiment 2, when neither a veridical cursor nor aiming target are provided, participants may lack a clear referent for their actual hand position (Synofzik, Thier, Leube, Schlotterbeck, & Lindner, 2010; Tsay, Kim, et al., 2021; Tsay, Kim, Haith, & Ivry, 2022). Here the system may not be able to fully dismiss the irrelevant cursor/target pair, resulting in a modest degree of adaptation near the irrelevant cursor/target location.

Second, the characteristics of the irrelevant visual errors may also modulate the degree to which this information is ignored. In Experiment 1b, the irrelevant visual errors were large and introduced abruptly. This may have led them to be easily deemed irrelevant, being outside the typical range of motor noise (Wei & Körding, 2009), and thus, not sufficient to elicit implicit adaptation. In contrast, in Experiment 2 the visual errors were small and were introduced more gradually, features that have made them be deemed as more relevant by the adaptation system (Ingram et al., 2000; Kagerer, Contreras-Vidal, & Stelmach, 1997). Future studies are required to systematically examine the characteristics that influence how the nervous system evaluates error relevance, and in turn, how this perceived relevance impacts the extent of implicit adaptation (e.g., (Makino, Hayashi, & Nozaki, 2023)).

### Limitations and future directions

A limitation of our study is that we did not control or monitor fixation. Since it is likely that participants directed their gaze at the relevant target (Neggers & Bekkering, 2000), one might argue that the dominance of the relevant cursor in driving adaptation could reflect the fact that distractor information is presented at more peripheral locations(s). We think that a fixation-based argument is unlikely to account for the effects observed here. First, the distractor cursors attenuated adaptation in all conditions, including in Experiment 2 where the relevant and distractor cursors were 90° apart. Thus, at least for the angular separations employed here, there is no obvious relationship between distance and attenuation. Second, prior work (Rand & Rentsch, 2015) has shown that magnitude of implicit adaptation is similar if participants are required to maintain fixation on the start position, target, or allowed to gaze freely.

Although the present work makes clear that adaptation is impacted when operating in a complex environment, one that is more akin to our natural environments, the processing stage at which this interaction occurs is unclear. The present work suggests the operation of a selection process that constrains the sensory prediction error, with the strength of this signal attenuated in the presence of task-irrelevant information. Alternatively, the presence of uncertainty may operate as some sort of gain on adaptation, either in terms of the strength of the sensory prediction or weight given to the sensory prediction error. An important question for future research is to elucidate the processes involved in the differential weighting of relevant and irrelevant sensory feedback, and specify how this information impacts adaptation.

## ACKNOWLEDGEMENTS

This work was supported by grants R35 NS116883, R01 NS105839, and R01 NS1058389 from the National Institutes of Health (NIH).

## Notes

### Competing Interest Statement

The authors have declared no competing interest.

### Summary of Updates

Author list updated. Modifications to main text to address reviewer comments.

